# Distinct kinetics and mechanisms of microbial inactivation of enteric virus revealed by capsid and genome Integrity

**DOI:** 10.64898/2026.09.23.753694

**Authors:** Josephine Meibom, L Daniela Morales, Htet Kyi Wynn, Sujin Shin, Tamar Kohn

**Affiliations:** Laboratory of Environmental Virology, School of Architecture, Civil and Environmental Engineering, École Polytechnique Fédérale de Lausanne (EPFL), Station 2, 1015 Lausanne, Switzerland

**Author notes:** To whom correspondence should be addressed: Tamar Kohn.

## Abstract

Enteric viruses are important contaminants of surface waters and a significant burden on public health. In aquatic ecosystems, the stability of these pathogens is differentially impacted by abiotic and biotic stressors, which can exert inactivating effects. Here, we investigated the fate of two enteroviruses, echovirus 11 (E11) and coxsackievirus B5 (CVB5), and one adenovirus, human adenovirus 2 (HAdV2), in lakewater and explored the mechanisms underlying their microbial inactivation. By combining infectivity assays, genome quantification, and capsid integrity analysis, we characterized virus-specific inactivation kinetics and mechanisms and examined the relationship between infectivity loss and capsid structural integrity. We observed rapid, intermediate, and negligible decay for HAdV2, E11, and CVB5, respectively, and confirmed that microbial proteases contribute to their inactivation. In addition, we revealed that loss of capsid structural integrity drives E11 inactivation, but not HAdV2 inactivation. Genome decay did not consistently correlate with loss of infectivity, highlighting the limitations of genome-based detection for assessing the presence of infectious viruses. These findings provide new insights into the mechanisms governing virus inactivation in aquatic environments and emphasize the importance of understanding viral fate when interpreting molecular detection data to assess virus-associated microbial risks.

**Importance:** Enteric viruses are common contaminants of surface waters and pose important risks to human health. However, the stability of these viruses in water varies widely and is influenced by many environmental factors. In this study, we investigated how microbial activity causes the inactivation of three enteric viruses in lakewater. We found substantial differences in virus stability and showed that microbial enzymes that break down proteins contribute to virus inactivation. We additionally found that virus structural damage is associated with the inactivation of one virus, but not of another, and that quantification of virus genetic material does not reliably indicate levels of infectious virus particles. These findings highlight the importance of understanding how viruses behave in surface waters and can help inform approaches for monitoring viral contamination and assessing potential risks to human health.

## Introduction

Enteroviruses and adenoviruses are two important groups of enteric viruses responsible for a large number of annual infections worldwide^1–3^. Enteroviruses are positive-sense single-stranded RNA viruses with small icosahedral capsids (approximately 30 nm in diameter) composed of repeating units of four structural proteins (viral proteins (VP) 1-4)^4,5^. Adenoviruses possess larger icosahedral capsids (approximately 90 nm in diameter) consisting of repeating units of three major (hexon, penton base, and fiber) and minor structural proteins enclosing a double-stranded DNA genome^6,7^. The fiber proteins protrude from the vertices of the adenovirus capsid and mediate host cell receptor attachment^8^, an important structural feature not present in enteroviruses.

Both virus groups are frequently detected in surface waters^9–12^, and thereby pose a risk to public health. For example, enteroviruses have been reported in 75% of surface water samples in the Netherlands^10^ while adenoviruses have been detected in 36.4% of recreational marine and freshwater sites across Europe^11^. Moreover, wide ranges of environmental stabilities have been reported for these pathogens^13^, with substantial stability differences also observed among closely rel ated virus types^14–16^. Such differences suggest that virus-specific inactivation mechanisms are critical in determining virus fate in surface waters. Although abiotic factors such as temperature and sunlight are known to contribute to both enterovirus and adenovirus decay^13^, the contribution of microbial mechanisms to virus inactivation and how they differentially impact these pathogens remain less clearly defined.

Aquatic microbial communities produce proteolytic enzymes that can contribute to virus inactivation^15–18^. Proteolytic degradation of the viral capsid may compromise structural integrity and lead to loss of infectivity^19–21^, yet the extent to which this mechanism governs virus decay and how it differs across virus types remain poorly understood. In particular, it is unknown whether microbial virus inactivation leads to capsid degradation to an extent that allows environmental nucleases to access and degrade the viral genome, as suggested for the enterovirus echovirus 12^14^, or whether viruses can lose infectivity while retaining structurally intact capsids that protect the viral genome. This distinction is critical for assessing the health risks associated with enteric viruses in surface waters as these pathogens may be detected using cell culture-based assays, which quantify infectious viral particles, or by molecular methods, which quantify viral genome copies. While molecular methods are rapid, sensitive, and virus-specific, they cannot distinguish infectious from non-infectious viral particles when the virus genome remains intact upon inactivation^22,23^. Understanding the mechanisms of virus inactivation is therefore critical for interpreting genome-based quantification and assessing how this reflects the infectious virus load in surface waters.

Herein, we investigated the microbial inactivation of two enteroviruses, echovirus 11 (E11) and coxsackievirus B5 (CVB5), and one adenovirus, human adenovirus 2 (HAdV2), in lakewater. By linking infectivity assays, genome quantification, and capsid integrity analysis, we characterized virus-specific inactivation kinetics and mechanisms and examined the relationship between infectivity loss and capsid structural integrity. Collectively, these findings provide new insights into the mechanisms governing virus inactivation in aquatic environments and highlight the importance of understanding viral fate when interpreting molecular detection data to assess virus-associated microbial risks.

## Materials and Methods

### Experimental approach

The stability of two enteroviruses, echovirus 11 (E11) and coxsackievirus B5 (CVB5), and one adenovirus, human adenovirus 2 (HAdV2), in lakewater was assessed by incubating these pathogens in Lake Geneva surface water samples sampled in three different times and monitoring their infectivity over time (**Figure 1.A**). In one of these lakewater samples, the mechanisms of microbial virus inactivation were further investigated through (i) analysis of viral genome degradation by digital PCR (dPCR), (ii) indirect assessment of virus capsid integrity by nuclease treatment, and (iii) direct assessment of enterovirus viral protein 1 (VP1) stability and HAdV2 fiber protein function by Western blot and host cell attachment, respectively (**Figure 1.B**). Detailed experimental procedures are described in the following sections.

**Figure 1.**
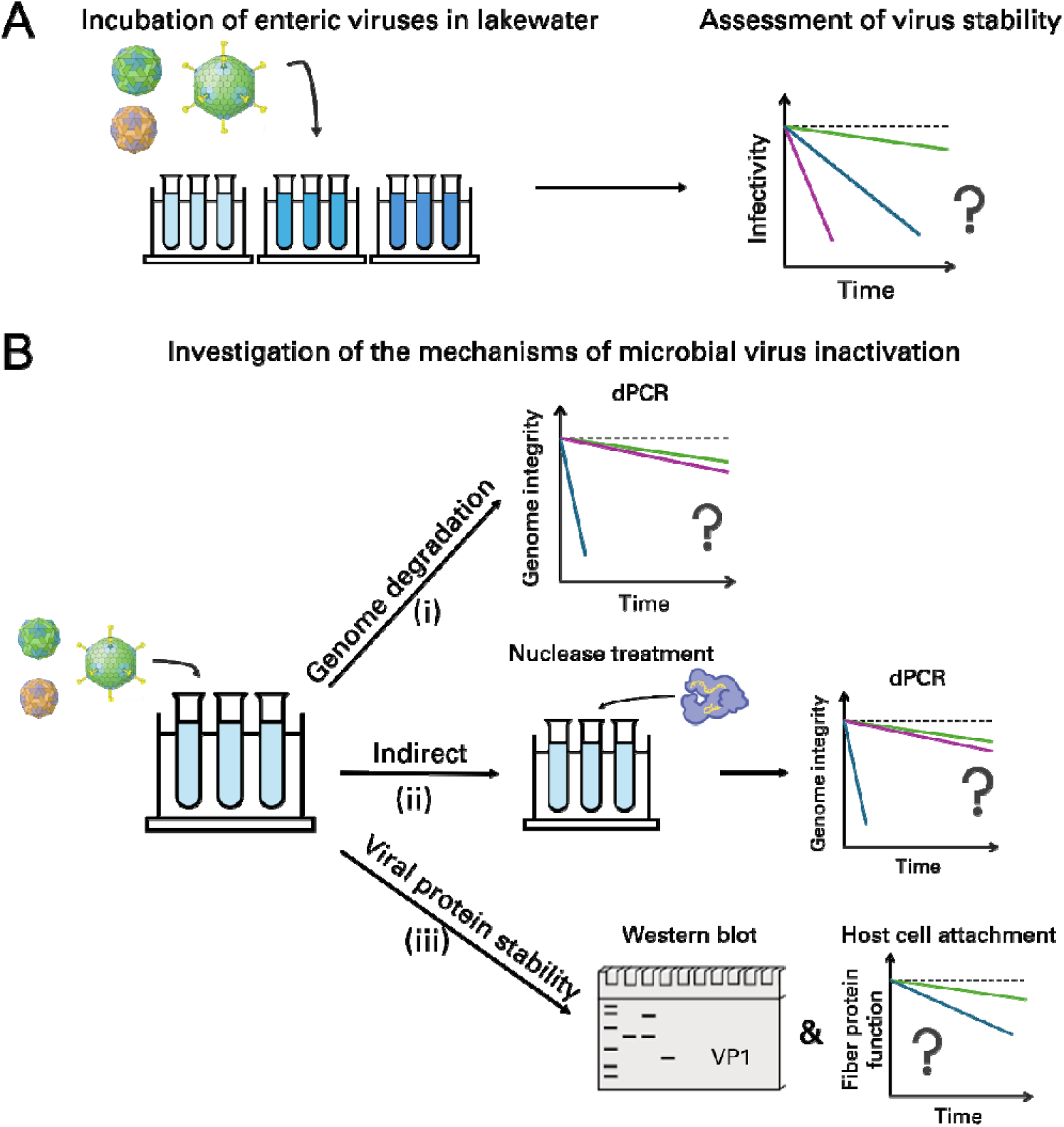
Experimental approach used in this study. (**A**) The stability of enteric viruses in lakewater was assessed by incubating the viruses in three Lake Geneva surface water samples and monitoring their infectivity over time. (**B**) Further characterization of the mechanisms of microbial virus inactivation was performed in one lakewater sample by (i) analysis of viral genome degradation, (ii) indirect assessment of virus capsid integrity, and (iii) direct assessment of enterovirus viral protein 1 (VP1) stability and HAdV2 fiber protein function.

### Virus propagation and purification

E11 (propagated from Gregory strain ATCC VR737), CVB5 (propagated from an environmental i solate, GenBank accession number MG845891), and HAdV2 (propagated from a stock kindly provided by Rosina Girones, University of Barcelona, Spain) stocks were prepared by infecting sub-confluent layers of rhabdomyosarcoma cells (RD, ATCC CCL-136), buffalo green monkey kidney cells (BGMK, kindly provided by Spiez Laboratory, Switzerland), or A459 cells (kindly provided by Rosina Girones from the University of Barcelona, Spain), respectively, in T-75 culture flasks. The cells were grown and maintained at 37°C and 5% CO_2_ with Dulbecco’s Modified Eagle Medium (DMEM, 41966, Gibco) or Minimum Essential Medium (MEM, A41922) supplemented with 1% penicillin-streptomycin (15140, Gibco) and 10% (growth) or 2% (maintenance) heat-inactivated fetal bovine serum (FBS, A5256701, Gibco). DMEM was used for RD and A549 cells while MEM was used for BGMK cells. The viruses were released from the cells three days post infection by freeze-thawing the culture flasks three times, and cell debris was removed by centrifugation at 1’100x *g* for 5 minutes. Virus stocks were prepared by buffer-exchange into phosphate-buffered saline (PBS, 18912014, Gibco) using Amicon Ultra-15 centrifugal filters (UFC910024, Merck Millipore). All virus stocks were tested for the presence of aggregates and were found to be monodisperse (**Supplementary Figure S1**). Details regarding virus size measurements can be found in the Supplementary Methods. HAdV2, a respiratory virus, is used here as a proxy for the enteric adenoviruses (HAdV40 and HAdV41) which are more difficult to culture and enumerate^24^.

High purity virus stocks for Western blot analysis of E11 and CVB5 were prepared by pelleting the cell culture supernatant through a 5 mL 20% sucrose cushion (38.5 mL ultracentrifuge tube: 344058, Beckman Coulter) at 150’000x *g* and 4°C for 3 hours in a SW 32 Ti S/N Swinging Bucket rotor (22U6711, Beckman Coulter). The obtained pellets were soaked overnight at 4°C in 50 µL PBS and then fully resuspended by pipetting up and down. The resuspended pellets were separated throughout a 15% – 50% sucrose gradient (17 mL ultracentrifuge tube: 344061, Beckman Coulter) at 150’000x *g* and 4°C for 3 hours in a SW 32 Ti S/N Swinging Bucket rotor (17U5130, Beckman Coulter). The sucrose gradient was prepared by layering 2 mL of 15%, 20%, 25%, 30%, 35%, and 40% and 4 mL of 50% sucrose solutions and resting overnight at 4°C to allow formation of a continuous gradient. The collected viral fractions were combined and buffer-exchanged into PBS using Amicon Ultra-15 centrifugal filters to yield high-purity virus stocks.

### Enumeration of virus infectivity

Infectious virions were enumerated using a most probable number (MPN) infectivity assay as described previously^25^. Briefly, RD, BGMK, or A549 cells were grown to 95% confluence in 96-well plates (Greiner CELLSTART, Sigma Aldrich). Then, 20 µL of each sample (in five replicates) were 10-fold serially diluted in maintenance medium and added to the cells. The plates were incubated at 37°C and 5% CO_2_ for 5 days (E11 and CVB5) or 7 days (HAdV2) after which time each well was visually examined for cytopathic effect (CPE) using an inverted microscope. The infectious virus titer was determined as the MPN of cytopathic units per milliliter (MPNCU/mL) by converting the number of wells positive for CPE using the R package *MPN*^26^. The assay limit of quantification (LOQ) was considered as one positive well (of five replicates) at the lowest dilution (10-fold diluted) and corresponded to 90.4 MPNCU/mL. Values below the LOQ were set to LOQ/√2, according to the recommendations of Hornung and Reed^27^, and are indicated by open symbols in all plots. The coefficient of variation of enumeration was determined to be 40-50% based on replicate titration of E11 (n = 12), CVB5 (n = 9), and HAdV2 (n=9). The titers of the virus stocks were 10^9^ or 10^11^ MPNCU/mL for E11, 10^9^ MPNCU/mL for CVB5, and 10^9^ MPNCU/mL for HAdV2. The titers of the high purity virus stocks were 10^8^ MPNCU/mL for E11 and 10^9^ MPNCU/mL for CVB5.

### Enumeration of virus genome copies

E11 and CVB5 genomes were enumerated using a reverse transcription digital PCR (RT-dPCR) assay as described previously^28^. Briefly, viral RNA was extracted from 140 μL sample using the QIAamp viral RNA Mini Kit (52906, Qiagen) according to the manufacturer’s instructions. The viral RNA was eluted in 60 μL elution buffer and stored at –20°C until quantification. Details regarding the enterovirus RT-dPCR assay can be found in the Supplementary Methods. Similarly, HAdV2 genomes were enumerated using a digital PCR (dPCR) assay. Briefly, viral DNA was extracted from 200 μL sample using the Maxwell RSC Blood DNA Kit (AS1520, Promega) and the Maxwell RSC Instrument (AS4500, Promega) according to the manufacturers’ instructions. The viral DNA was eluted in 60 μL elution buffer and stored at –20°C until quantification. Details regarding the HAdV2 dPCR assay can be found in the Supplementary Methods.

Quantities were expressed as genome copies per mL of sample (GC/mL) and genome decay was expressed as Log_10_(N/N_0_) where N is the quantified genome copies at time *t* (hours) and N_0_ is the quantified genome copies at time 0 hours. The assay limit of quantification (LOQ) was considered as two positive partitions, corresponding to a sample concentration of 1.04*10^3^ GC/mL. Data points below the LOQ were set to LOQ/√2, according to the recommendations of Hornung and Reed,^27^ and are indicated by open symbols in all plots.

### Lakewater sampling and processing

Surface lakewater was collected from Lake Geneva on April 9^th^, 2024, July 23^rd^, 2024, and March 2^nd^, 2026, from the shore in Saint-Sulpice, Switzerland. Immediately after collection, the lakewater samples were brought to the laboratory and vacuum-filtered through a 0.8 μm MCE membrane filter (AAWP04700, MF-Millipore) to remove eukaryotes while retaining bacteria in the lakewater^29^. The resulting filtrate was termed microbially active lakewater. Sterile lakewater was obtained by autoclaving the filtered lakewater at 121°C for 15 minutes. Both microbially active and sterile lakewater samples were stored at 4°C overnight prior to virus inactivation experiments.

### Virus inactivation experiments in lakewater

Inactivation experiments with E11, CVB5, and HAdV2 were conducted in all lakewater samples. Briefly, 1-10 mL of microbially active and sterile lakewater samples were separately spiked with virus to an initial virus titer of approximately 10^7^ MPNCU/mL. Aliquots of 150 μL were taken at timepoints across 54 hours and stored at –20°C until enumeration of virus infectivity. In March 2026, additional aliquots were sampled at each timepoint for enumeration of viral genome copies, indirect assessment of capsid integrity, and assessment of HAdV2 fiber protein function. Each experiment was conducted in triplicate, and the samples were maintained shaking gently at room temperature (KS 501 digital shaker, IKA) throughout the experiment. Virus inactivation rate constants (*k*, h^-1^) were determined by least square fit of inactivation data pooled across triplicate experiments to a first-order decay model (**Equation 1**):

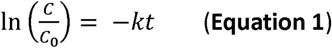

where *C* is the infectious virus titer at time *t* (h) and *C_0_* is in initial infectious virus titer. Data points below the LOQ were not considered for the determination of rate constants. Values of *k* are reported with 95% confidence intervals (CI) associated with the regression slope (**Supplementary Table S3**).

### Indirect assessment of virus capsid integrity

Virus capsid integrity was assessed indirectly by treating samples with nucleases (RNase for enteroviruses, DNase for adenovirus) prior to enumeration of viral genome copies. This analysis is based on the hypothesis that an intact capsid protects the viral genome from degradation by nucleases^30^, whereas capsid decay exposes the genome to nuclease-mediated degradation. The experiment was conducted in lakewater sampled in March 2026, following the protocol described previously^31^. Briefly, 30 μL aliquots of E11 or CVB5 spiked into microbially active and sterile lakewater were taken at timepoints across 54 hours, immediately mixed with 3 μL RNase T1/A Mix (EN0551, Thermo Scientific) and 107 μL RNA digestion buffer (10 mM Tris-HCl and 1 mM EDTA pH 7.4 (93392, Supelco) supplemented with 50 mM NaCl), and incubated at 37°C for 30 minutes (water bath). Subsequently, the reaction was quenched by adding 1 μL SUPERase-In RNase Inhibitor (AM2694, Invitrogen) and incubating at room temperature for 20 minutes. Similarly, 43 μL aliquots of HAdV2-spiked microbially active and sterile lakewater were taken at timepoints across 54 hours, immediately mixed with 10 μL DNase I Solution (89836, Thermo Scientific) and 147 μL 1x DNA digestion buffer (supplied with the DNase I Solution), and incubated at 37°C for 30 minutes (water bath). Subsequently, the reaction was quenched by adding 10 μL EDTA 50 mM (89836, Thermo Scientific) and incubating at 65°C for 10 minutes. All samples were then stored at –20°C until enumeration of viral genome copies.

### Direct assessment of enterovirus viral protein 1 (VP1) stability

Enterovirus structural integrity was directly assessed through Western blotting targeting viral protein 1 (VP1) in lakewater sampled in March 2026. Briefly, samples were prepared by concentrating sucrose-purified E11 and CVB5 virus stocks using 10k MWCO PES Pierce Protein Concentrators (88513, Thermo Scientific) and adding the concentrated virus to microbially active lakewater, sterile lakewater, and PBS at a protein concentration of 0.02 – 0.05 mg/mL. Aliquots of 50 μL were taken at each timepoint, immediately mixed with 10 μL Laemmli SDS sample reducing buffer (6x, J61337-AC, Thermo Scientific), and boiled at 95°C for 15 minutes to denature the viral proteins. The samples (40 μL) were loaded onto Novex 4%-20% Tris-Glycine Plus WedgeWell protein gels (XP04200BOX, Invitrogen) and run at 140 V for 75 minutes. Separated viral proteins were transferred onto nitrocellulose membranes using the iBlot 2 Dry Blotting System (iBlot 2 Mini Transfer Stacks and iBlot 2 Gel Transfer Device, IB23002 and IB21001, Invitrogen) at 15 V for 15 minutes. Subsequently, the membrane surface was blocked with 5% skim milk (70166, Fluka) in 1x PBS-Tween 20 (28352, Thermo Scientific) for 30 minutes. VP1 was targeted with Rabbit Anti-Enterovirus Polyclonal Antibody (1:1000 dilution, PAB21467, The Native Antigen Company) and detected following conjugation with horseradish peroxidase labeled secondary antibody (Rabbit IgG HRP Linked Whole Ab (from Donkey), NA934, Cytiva) using the WesternBright ECL spray (K-12049-D50, Advansta) for blot development. Washing steps (twice with Tween 20 1x in PBS, 28352, Thermo Scientific) were included after antibody incubations to remove unbound antibody. The Fusion Fx imaging system (Vilber S.A.) was used for imaging with 10 seconds exposure time. In parallel, aliquots of 5 μL were taken across 54 hours, mixed with 495 μL PBS and stored at –20°C until enumeration of virus infectivity.

### Direct assessment of HAdV2 fiber protein function

HAdV2 fiber protein function was evaluated by assessing adenoviral host cell attachment capability, following a method adapted from Gall et al.^32,33^ The experiment was conducted in triplicate in lakewater sampled in March 2026. Briefly, 200 μL aliquots of HAdV2 spiked into microbially active and sterile lakewater were taken at timepoints across 54 hours and diluted 10-fold in DMEM supplemented with 1% penicillin-streptomycin and 2% FBS. Then, 2 mL of diluted sample were inoculated onto pre-cooled (4°C, 1 hour) A549 cell monolayers in 6-well plates and incubated at 4°C for 90 minutes under gentle shaking to allow virus binding to the host cells while minimizing virus internalization. Following incubation, the inoculum was removed by aspiration, and the A549 monolayers were washed twice with cold (4°C) PBS to remove unbound virus. Subsequently, the cells were trypsinized, collected, and subject to three freeze-thaw cycles. Cell debris was removed by centrifugation at 3’000x *g* for 5 minutes and viral DNA was extracted from the supernatant using the PureLink Viral RNA/DNA Mini Kit (12280050, ThermoFisher Scientific) according to the manufacturer’s instructions. Viral DNA was eluted in 50 μL elution buffer and stored at –20°C until quantification by long-range PCR followed by quantitative PCR (qPCR). Details regarding the PCR-qPCR assay targeting the adenovirus hexone gene can be found in the Supplementary Methods. A previously reported calibration curve of the HAdV2 amplicon^33^ was used for calculation of genome copies per mL of sample (GC/mL). Loss of fiber protein function was expressed as decay Log_10_(N/N_0_) of the HAdV2 amplicon, where N is the quantified genome copies at time *t* (hours) and N_0_ is the quantified genome copies at time 0 hours.

### Metalloprotease inhibitor experiment

The effect of the metalloprotease inhibitor GM6001 (CC1010, Sigma Aldrich) on virus inactivation was studied in July 2024. Briefly, GM6001 was added into microbially active lakewater at a concentration of 5 μM and incubated at room temperature for 30 minutes prior to performing virus inactivation experiments.

### Statistical analysis

Assessment of the correlation between HAdV2 inactivation and loss of fiber protein function was performed by calculating Pearson’s correlation coefficient (r) assuming a normal distribution. Statistical comparison between virus inactivation in lakewater with and without protease inhibitor was performed by paired t-test using a significance threshold of α = 0.05. All statistical analyses were conducted in GraphPad Prism v10.6.1 (GraphPad Software, USA).

## Results

### E11 and HAdV2 are microbially inactivated in lakewater while CVB5 remains mostly stable

To evaluate the stability of three enteric viruses in lakewater, we incubated E11, CVB5, and HAdV2 in microbially active Lake Geneva surface water samples collected in April 2024, July 2024, and March 2026, and monitored infectivity over time (**Figure 2**). We observed first-order decay of E11 in all samples, resulting in similar inactivation rate constants (*k*: 0.09 ± 0.01 h^-1^ to 0.11 ± 0.01 h^-1^, **Supplementary Table S3**). While CVB5 exhibited linear decay in July 2024 (*k*: 0.12 ± 0.01 h^-1^), this virus remained stable in samples collected in April 2024 and March 2026. We observed biphasic inactivation kinetics for HAdV2 in April 2024 and March 2026, with rapid initial linear decay over 30 hours (*k*: 0.29 ± 0.03 h^-1^ in April 2024, 0.25 ± 0.02 h^-1^ in March 2026) followed by stabilization of the virus titer. In contrast, HAdV2 exhibited linear decay following an initial 24 hour period of stability in July 2024, yielding an inactivation rate constant (*k*: 0.10 ± 0.01 h^-1^) similar to those of E11 and CVB5 in the same lakewater sample. Although virus decay was observed in some sterile samples, possibly caused by microbial contamination and regrowth throughout the incubation period, we consistently observed greater decay in microbially active lakewater. This confirmed the importance of microbial mechanisms for the inactivation of enteric viruses in lakewater.

**Figure 2.**
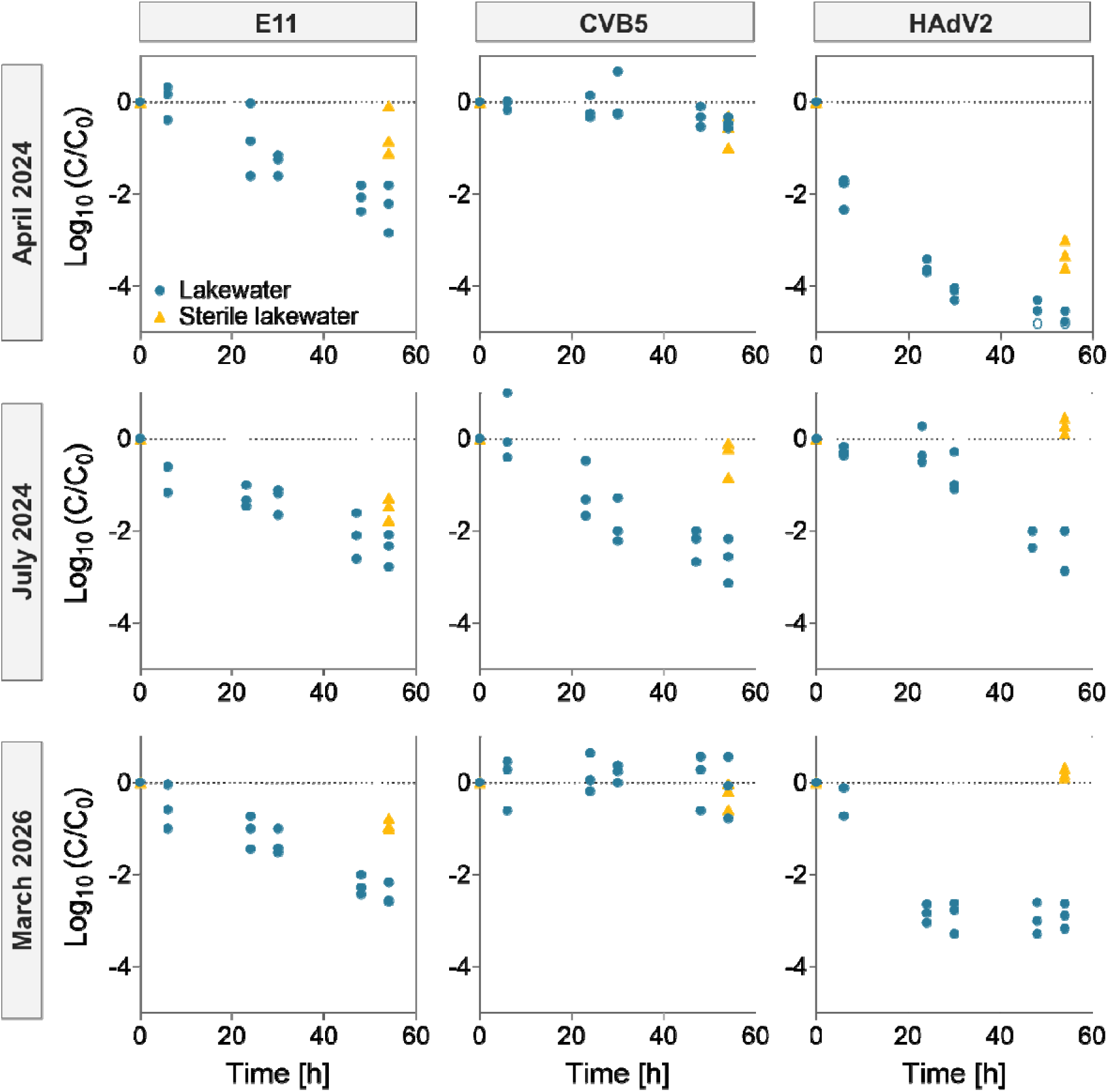
Inactivation of E11, CVB5, and HAdV2 in microbially active (blue circles) and sterile (orange triangles) Lake Geneva surface water collected in April 2024, July 2024, and March 2026. Individual data points of triplicate experiments are presented. Data points below the detection limit were set to a concentration of LOQ/√2 and indicated by an open symbol. Inactivation rate constants are given in **Supplementary Table S3.**

The inactivation rates observed in April 2024 and March 2026 (**Figure 2**) indicated that E11, CVB5, and HAdV2 possess markedly different stabilities in lakewater. In both samples, CVB5 remained stable throughout the incubation period while E11 displayed intermediate stability, consistent with previous observations^16^, and HAdV2 exhibited rapid initial decay followed by stabilization of the virus titer. In contrast, CVB5 and HAdV2 exhibited very different inactivation profiles in July 2024 (**Figure 2**). In this sample, CVB5 was not stable and HAdV2 displayed an initial period of stability followed by rapid inactivation. Overall, all three viruses exhibited similar inactivation rates (**Supplementary Table S3**), suggesting atypical antiviral activity in July 2024.

To confirm the contribution of proteases to virus inactivation, we determined the stability of E11, CVB5, and HAdV2 in lakewater in the presence of the metalloprotease inhibitor GM6001 (**Supplementary Figure S2**). Inhibition of metalloproteases was selected for assessment of proteolytic inactivation as this class of proteases is known to cause enterovirus decay^15^. We observed a significant reduction (> 0.9 Log_10_ units; p < 0.1) in the inactivation of E11 and CVB5 after 54 hours, in agreement with previous findings^16^. The effect was also measurable, though it was less significant for HAdV2 (0.8 Log_10_ units; p = 0.12). The importance of microbial, and specifically proteolytic, mechanisms for the inactivation of enteric viruses in lakewater was thus confirmed.

### Genome decay does not correlate with virus inactivation

To characterize the mechanism of microbial inactivation of enteric viruses in lakewater, we monitored genome decay by digital PCR (dPCR) in March 2026 (**Figure 3**, middle column). Proteolytic decay has been shown to cause genome release through degradation of the virus capsid^19^, thereby exposing the viral genome to nucleases present in the lakewater matrix. While dPCR quantifies only a short segment of the viral genome, nuclease-mediated viral genome degradation is assumed to affect the entire genome, including the segment targeted by dPCR. We therefore interpret any decay in the genome segment targeted by dPCR to reflect overall genome decay.

**Figure 3.**
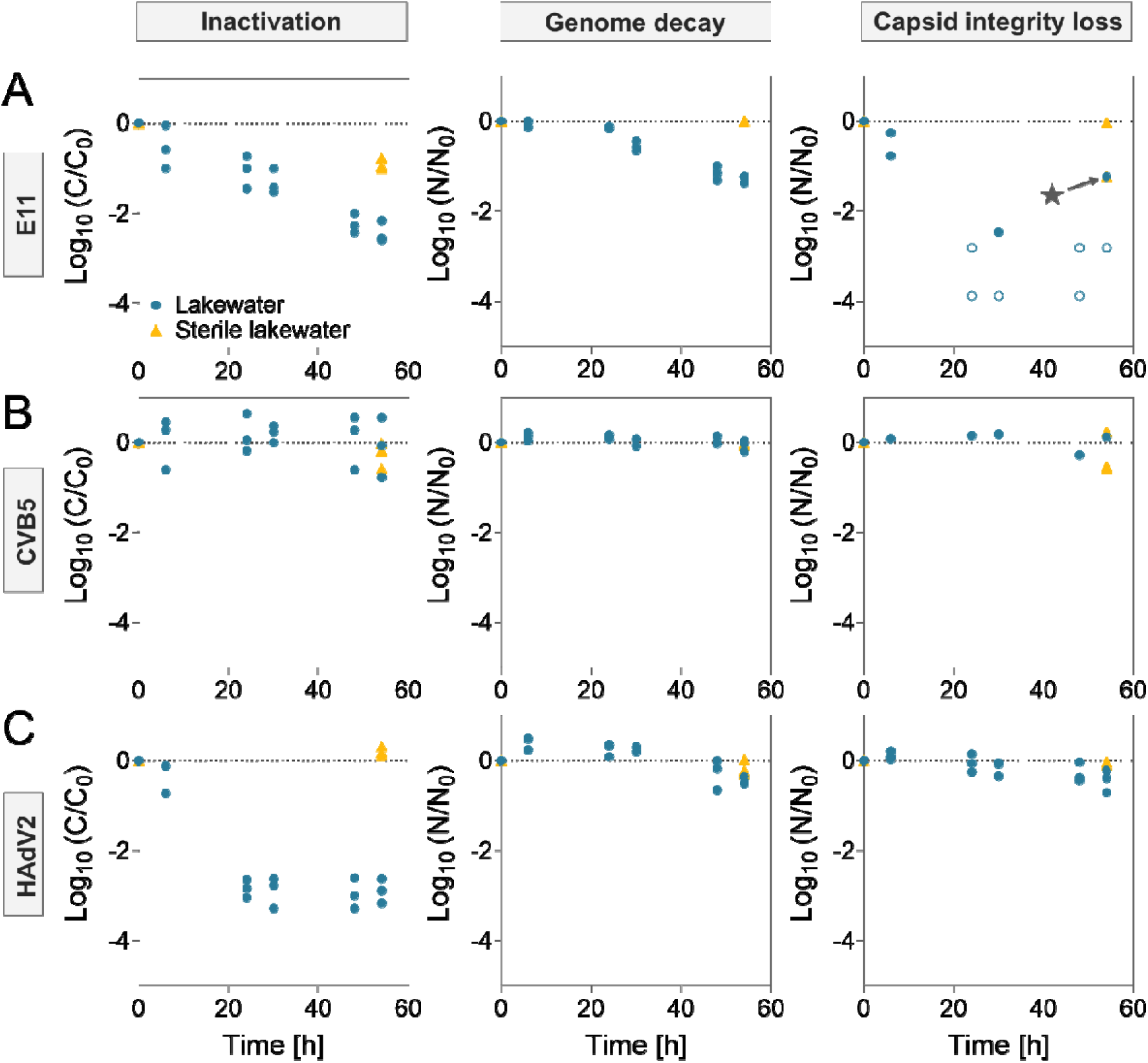
Comparison of inactivation, genome decay, and capsid integrity loss (genome decay after nuclease treatment) of **(A)** E11, **(B)** CVB5, and **(C)** HAdV2 in microbially active (blue circles) and sterile (orange triangles) Lake Geneva surface water collected in March 2026. Inactivation data (left column) were previously presented in Figure 2. Virus genome copies were quantified before (middle column) and after (right column) nuclease treatment. Individual data points of triplicate experiments are presented, except for nuclease-treated E11 and CVB5 samples for which two and one replicates are presented, respectively. Data points below the dPCR detection limit were set to a concentration of LOQ/√2 and indicated by an open symbol. Note that the different measurable ranges of genome decay result from differences in the starting concentrations of the replicates. The data point indicated by a star is an outlier and indicates likely contamination.

In general, we observed lower genome decay compared to loss of infectivity after 54 hours (**Figure 3**). We observed a 1.3 Log_10_ unit decrease in genome copies for E11 compared to a 2.4 Log_10_ reduction in infectivity (**Figure 3.A**). In contrast to inactivation, no genome decay was detected during the first 24 hours, suggesting either slow permeabilization of the E11 capsid or slow RNase activity in the lakewater. Genome copies remained stable for CVB5 throughout the incubation period, in agreement with the stability of this virus in lakewater (**Figure 3.B**). We observed no decay in genome copies for HAdV2 over 54 hours despite a 2.9 Log_10_ reduction in infectivity (**Figure 3.C**), suggesting no significant damage to the HAdV2 capsid and no genome release. These results reveal that genome decay does not consistently correlate with inactivation and indicate that viral capsids remain at least partially intact in lakewater.

### Loss ofcapsid structural integrity drives E11 inactivation but not HAdV2 inactivation

To further assess virus capsid integrity in lakewater, we treated the samples with nucleases prior to genome extraction and quantification by dPCR (**Figure 3**, right column) to enhance the possibly low nuclease activity in lakewater. A reduction in genome copies after nuclease treatment indicates loss of virus structural integrity as a permeable capsid allows degradation of the viral genome by exogenous nucleases^30^.

Unexpectedly, lower absolute genome copy numbers were measured after RNase treatment for E11 and CVB5 compared to samples without RNase treatment (**Supplementary Figure S3**) and not all replicates were quantifiable at early timepoints. These results were consistent across both diluted and undiluted RNA extracts (**Supplementary Figure S4**), indicating that PCR inhibition was not the cause of this discrepancy. Additionally, no differences in genome copy numbers before and after nuclease treatment were observed in preliminary control experiments for all three viruses (**Supplementary Figure S5**). Despite the unexplained low genome copy numbers in the RNase treated samples, trends in Log_10_(N/N_0_) genome decay (**Figure 3**, right column) could nevertheless be determined.

We observed rapid decay of E11 genome copies following RNase treatment (**Figure 3.A**), indicating loss of capsid integrity. In contrast to genome decay measured before RNase treatment, no lag phase in genome decay was observed after RNase treatment, suggesting that the E11 capsid initially retains sufficient integrity to protect the E11 genome in the absence of exogenous RNases. We observed stable CVB5 genome copies after RNase treatment (**Figure 3.B**), indicating preservation of capsid integrity and agreeing with the high stability of CVB5 in lakewater. Interestingly, genome copy numbers remained unchanged for HAdV2 before and after DNase treatment (**Figure 3.C**), suggesting that the HAdV2 capsid remains intact and protective, despite rapid inactivation. Nuclease treatment thus revealed important differences in capsid structural integrity among the three viruses in lakewater.

### Degradation of viral protein 1 drives enterovirus inactivation

To directly assess enterovirus capsid degradation during inactivation in lakewater, we performed Western blotting targeting viral protein 1 (VP1, **Figure 4**). VP1 (approximately 32 kDa), one of the four structural proteins that make up the enterovirus capsid, is the most surface exposed viral protein and is important in mediating host cell attachment^34^. To evaluate the relationships between VP1 degradation and enterovirus inactivation, we included samples incubated in phosphate-buffered saline (PBS) as controls and measured infectivity in all samples (**Supplementary Figure S6**).

**Figure 4.**
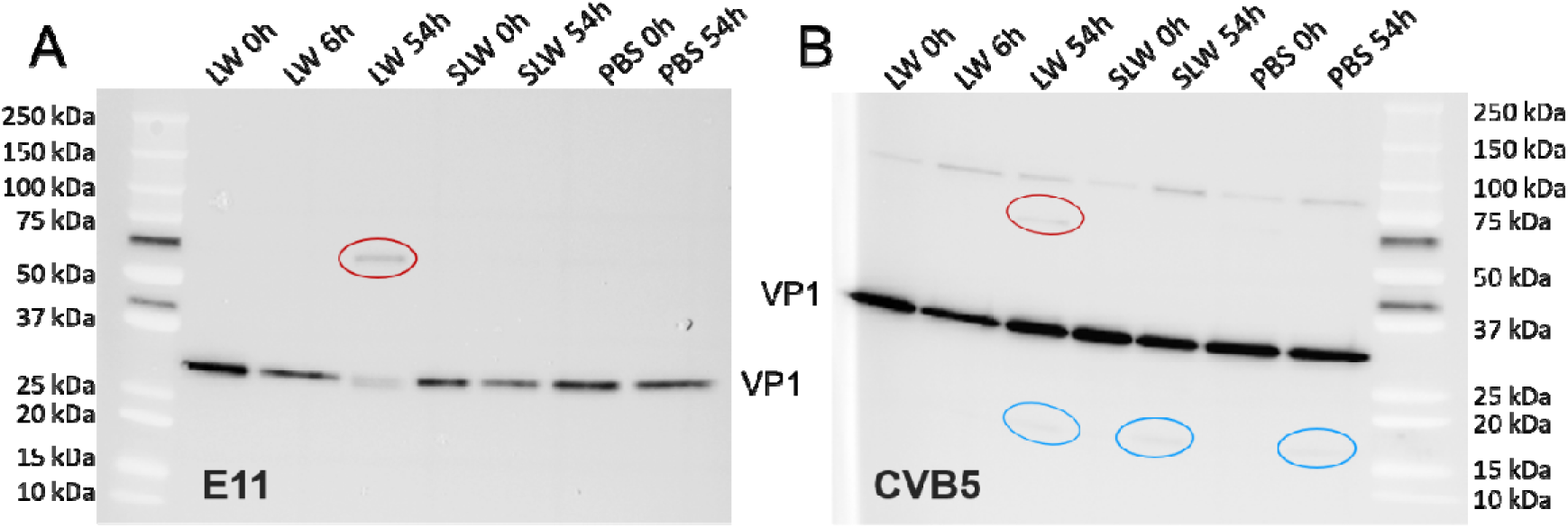
Western blot analysis of viral protein 1 (VP1) for **(A)** E11 and **(B)** CVB5 after incubation of the viruses in microbially active lakewater (LW), sterile lakewater (SLW), or phosphate-buffered saline (PBS). Degradation products and aggregates are indicated by blue and red circles, respectively.

We observed a reduction in the band corresponding to VP1 of E11 in microbially active lakewater after 54 hours (**Figure 4.A**), consistent with E11 inactivation (**Supplementary Figure S6.A**). At this timepoint, a higher molecular weight band (red circle) was also detected, suggesting VP1 aggregation or interaction with other (viral or lakewater) proteins. At 6 hours, VP1 remained stable and no aggregation was observed, in agreement with the stable genome copies measured for E11 before RNase treatment at this timepoint (**Figure 3.A**, middle column) and confirming the initial retention of E11 capsid integrity despite inactivation (**Figure 3.A**, left column). No degradation or aggregation of VP1 was observed in sterile lakewater or in PBS (**Figure 4.A**), in agreement with the stability of E11 in these matrices. We observed a highly stable VP1 band for CVB5 in lakewater across 54 hours (**Figure 4.B**), consistent with the measured infectious stability (**Supplementary Figure S6.B**). At 54 hours, faint bands at higher (red circle) and lower (blue circle) molecular weights were observed, suggesting limited protein aggregation or interaction, and degradation of VP1, respectively. The lower molecular weight band was also observed in sterile lakewater and PBS after 54 hours, although no degradation of VP1 was observed. A faint band at approximately 90 kDa was detected in all CVB5 samples, possibly indicative of VP1 aggregates formed during preparation of the virus stock or insufficient denaturing conditions. Overall, these results confirm that enterovirus inactivation in lakewater is associated with loss of capsid structural integrity.

### Adenovirus inactivation is associated with loss of fiber protein function

A Western blot assay targeting the fiber protein of adenovirus, which protrudes from the capsid and mediates binding to the host cell coxsackie-adenovirus receptor (CAR)^8^, proved unsuccessful. Instead, we assessed HAdV2 attachment to host cells to determine whether fiber protein degradation contributes to HAdV2 inactivation in lakewater (**Figure 5**).

**Figure 5.**
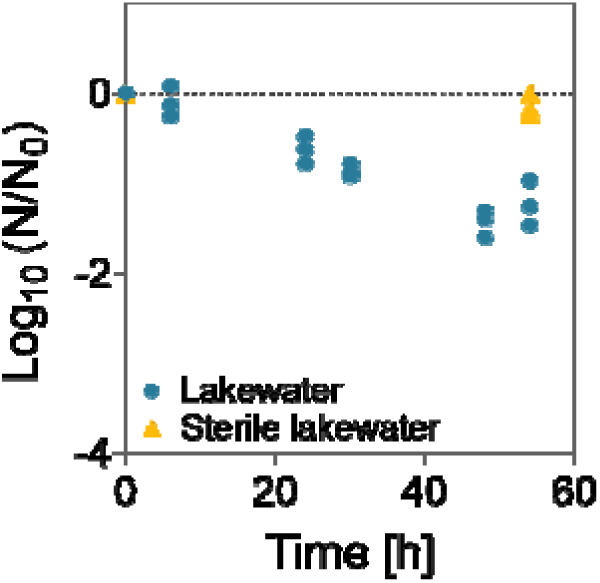
Loss of HAdV2 fiber protein function in microbially active (blue circles) and sterile (orange triangles) Lake Geneva surface water collected in March 2026. Individual data points of triplicate experiments are presented.

We observed loss of HAdV2 host cell attachment capacity over 54 hours (**Figure 5**, **k**: 0.06 ± 0.01 h^-1^), indicating loss of fiber protein function. While loss of HAdV2 fiber protein function was lower than the observed loss of infectivity (**Figure 3.C**), a positive correlation was found between these functions (Pearson correlation coefficient r = 0.89) supporting a role for fiber protein degradation in the inactivation of HAdV2 in lakewater.

## Discussion

We evaluated the stability of two enterovirus types (E11 and CVB5) and one adenovirus type (HAdV2) in lakewater. These viruses exhibited markedly different stabilities in microbially active lakewater (**Figure 2**), with rapid, intermediate, and negligible decay generally observed for HAdV2, E11, and CVB5, respectively. This trend was reflected in two of the three lakewater samples studied here (April 2024 and March 2026) and was consistent with previous observations for E11 and CVB5 in additional samples^16^. The different inactivation rates observed for each virus mirrors reported differences in susceptibility to disinfectants such as ozone^35^ and chlorine^36,37^ that act through oxidative mechanisms that disrupt the viral capsid^38^. We found that microbial proteases contribute to the inactivation of these pathogens in lakewater (**Supplementary Figure S2**), suggesting that cleavage of the proteinaceous capsid is a key mechanism driving microbially-mediated enteric virus decay in lakewater.

In contrast to April 2024 and March 2026, lakewater collected in July 2024 yielded substantially different results (**Figure 2**). In this sample, we observed similar inactivation rates for all three viruses (**Supplementary Table S3**), with notable decay of the typically stable CVB5 and slower overall decay of HAdV2 following an initial period of stability. In this sample, metalloproteases contributed to the decay of all three viruses (**Supplementary Figure S2**) and the proteolytic fingerprint closely resembled those of other Lake Geneva samples^39^, indicating that the unusual inactivation rates observed in July 2024 were not caused by a fundamentally different protease pool. The proteolytic nature and extent of virus inactivation in July 2024 may be associated with the presence of lakewater compounds capable of increasing the susceptibility of the virus to proteolytic attack. For example, epsilon-poly-L-lysine, an antimicrobial biopolymer produced by bacteria, has been shown to promote the proteolytic inactivation of *Calciviridae* by inducing capsid conformational changes^40^. Alternatively, other antiviral microbial compounds such as peptides^41,42^, extracellular polymeric substances^43,44^, or surfactants^45^, may have contributed to virus decay in this sample. Such alternative antiviral microbial mechanisms in lakewater are an important avenue for future investigation.

The enhanced stability of E11 in the presence of a metalloprotease inhibitor (**Supplementary Figure S2**) confirmed previous findings^15,16^ that the microbial inactivation of E11 in lakewater is mediated through the action of proteases. Previous studies have reported that proteases drive enterovirus decay through cleavage of the proteinaceous capsid^17,19^. We confirmed this mechanism for E11 inactivation by observing degradation of VP1, one of the four structural proteins of enteroviruses, by Western blotting (**Figure 4.A**) and by measuring genome decay, particularly following RNase treatment (**Figure 3.A**). Although loss of structural integrity thus drives E11 inactivation, capsid degradation is not immediate. We observed slower genome decay compared to inactivation over 54 hours (**Figure 3.A**), with genome decay occurring following a 24 hour delay. This suggests that the E11 capsid remains at least partly intact for a short period before being degraded, sufficient to initially protect the viral genome. This lag phase was confirmed by the observed stability of VP1 after 6 hours incubation (**Figure 4.A**) and has previously been reported for E11 in Lake Geneva^28^. Together, these results indicate that the E11 capsid initially retains structural integrity before undergoing progressive degradation in lakewater.

We found that metalloprotease inhibitors also reduce HAdV2 decay in lakewater (**Supplementary Figure S2**), suggesting that proteolytic degradation mediates HAdV2 inactivation. However, we observed stable HAdV2 genome copies throughout the incubation period (**Figure 3.C**), indicating that the main capsid structure remains intact. High HAdV2 structural stability is not unexpected as double-stranded DNA viruses often exhibit high internal pressure and mechanical stability due to the difficulty of compressing a double stranded, rather than a single-stranded, genome^46^. For example, HAdV2 can withstand forces of 3.3 nN^47^, while the single-stranded RNA enterovirus coxsackievirus A6 can only tolerate 1.12 nN^48^. A highly robust HAdV2 capsid body may therefore be resistant to proteolytic attack. In contrast, the fiber proteins contain the host cell binding sites that interact with the coxsackievirus and adenovirus receptor (CAR)^8^ implying that even minor damage to the CAR binding site could inhibit attachment and prevent infection. Additionally, the fiber proteins protrude from the vertices of the HAdV2 capsid and are thus highly accessible to proteases. The observed positive correlation between loss of fiber protein function and virus decay indicates that fiber protein degradation plays a role in HAdV2 inactivation. However, the slower loss of fiber protein function relative to virus inactivation suggests that other factors may also play a role in HAdV2 decay. For example, a conserved RGD motif in the penton base serves to bind the α_v_β_3_ and α_v_β_5_ integrin host cell co-receptor after CAR binding, an important step in virus internalization^49^. Degradation or structural disruption of the penton base protein may thus additionally contribute to HAdV2 decay. The role of these structural proteins in the proteolytic inactivation of HAdV2 in lakewater warrants further investigation.

The biphasic inactivation profile observed for HAdV2 in lakewater contrasted with the first-order decay profile of E11 (**Figure 2**). Such biphasic decay has previously been reported for HAdV2 incubated with the eukaryotic fraction of lakewater^29^ and during viral treatment with free chlorine^50,51^ and chlorine dioxide^52^, where it was attributed to the presence of viral aggregates. As no aggregates were observed in the HAdV2 stock used here (**Supplementary Figure S1**), such an effect cannot explain the observed tailing in lakewater. However, other factors may play a role. For example, selective degradation of the fiber proteins by microbial proteases may release small cleavage products forming a protective layer on the virus surface, as observed during chlorine dioxide treatment of the bacteriophage MS2^53^. Alternatively, incomplete cleavage of the twelve fiber proteins could lower the probability of infectivity while not entirely eliminating it.

Overall, we found that genome decay did not consistently correlate with loss of infectivity (**Figure 3**). Although the E11 genome exhibited linear decay following a 24 hour delay, total genome loss remained smaller than virus inactivation over the full incubation period. Thus, quantification of the E11 genome may overestimate infectivity unless this lag phase is accounted for. In contrast, HAdV2 genome copies remained stable throughout time despite rapid loss of infectivity, revealing that quantification of the HAdV2 genome does not reflect infectivity. While lower inactivation rate constants in surface water have typically been reported for enteroviruses when using molecular rather than cell culture methods, this is not usually the case for adenoviruses^13^. The divergence from the literature may be due to the contribution of other decay mechanisms, such as solar inactivation, to adenovirus inactivation in surface water which were not considered here. Additionally, while we expect full genome degradation by nucleases upon genome release from degraded capsids, including the genome region targeted by dPCR in this study, this assumption was not verified. Thus, the persistence of detectable HAdV2 genomes may reflect the persistence of genome fragments rather than intact, infectious virus, highlighting an important limitation of molecular methods for assessing viral persistence.

Despite the rapid microbial inactivation of HAdV2 in lakewater, literature data shows an overall greater stability of adenoviruses rather than enteroviruses in surface water^13^. This discrepancy may be due to other environmental factors not considered herein that contribute to the overall inactivation of enteric viruses. In particular, sunlight and temperature play key roles in the decay of both enteroviruses and adenoviruses, with adenoviruses exhibiting greater resistance to solar inactivation compared to enteroviruses^13,54^. To place our observations in this broader context, we estimated solar inactivation rate constants in Lake Geneva under summer conditions^29^ using published data on the solar inactivation of enteric viruses^33,55,56^ (**Table 1**). This analysis confirmed the slower decay of HAdV2 under solar irradiation and provides a potential explanation for the greater overall environmental stability of adenoviruses relative to enteroviruses^13^. Taken together, these results indicate that microbially mediated inactivation represents one of several factors governing the environmental fate of enteric viruses. Although often overlooked, this mechanism can vary substantially across virus groups and species and may therefore contribute significantly to differences in overall virus stability in natural waters.

**Table 1.** Microbial inactivation rate constants (*k_microbial_*) measured herein and estimated solar inactivation rate constants (*k_solar_*) for E11, CVB5, and HAdV2. Solar inactivation rate constants are based on published rate constants^33,55,56^ and adapted to reflect inactivation by solar UVB (280 to 320 nm) in Lake Geneva surface lakewater on a summer day, as described elsewhere^29^.

| Virus | $k_{microbial}$ [h<sup>-1</sup>] | $k_{solar}$ [h<sup>-1</sup>] |
| --- | --- | --- |
| **E11**<sup>56</sup> | 0.09 – 0.11 | 0.3 – 0.7 |
| **CVB5**<sup>55</sup> | 0.00 – 0.12 | 1.2 |
| **HAdV2**<sup>33</sup> | 0.10 – 0.29 | 0.1 |

Collectively, these findings highlight the broad range of enteric virus stability towards microbial decay in lakewater and provide mechanistic insight into protease-mediated virus decay. By combining infectivity assays, genome quantification, and capsid integrity analysis, we show that capsid degradation drives E11 inactivation whereas HAdV2 decay likely results from proteolytic damage to protruding fiber proteins rather than to the capsid body. These results may inform the selection of virus detection methods, infectivity versus genome quantification, for assessing viral risk in surface waters.

## Supporting information

Meibom-et-al 2026 Supplemental

## Acknowledgements

This work was funded by the Swiss National Science Foundation (grant no. 310030 215226). We thank Vidhi Dholakia for assisting with DLS measurements.

Conceptualization: J.M. and T.K. Investigation: J.M., L.D.M., H.K.W., and S.S. Methodology: J.M., L.D.M., S.S., and T.K. Data curation: J.M. Formal analysis: J.M. Visualization: J.M. Resources: T.K. Funding acquisition: T.K. Supervision: T.K. Writing – original draft: J.M. and T.K. Writing – review and editing: J.M., L.D.M, S.S., and T.K.

## Data Availability

All data are available on Zenodo under link: https://doi.org/10.5281/zenodo.22893716.

## References

(1) CDC. Non-Polio Enterovirus Outbreaks. Non-Polio Enterovirus. https://www.cdc.gov/non-polio-enterovirus/outbreak-surveillance/index.html (accessed 2025-10-17)

(2) MacNeil, K. M.; Dodge, M. J.; Evans, A. M.; Tessier, T. M.; Weinberg, J. B.; Mymryk, J. S. Adenoviruses in Medicine: Innocuous Pathogen, Predator, or Partner. Trends in Molecular Medicine 2023, 29 (1), 4–19. 10.1016/j.molmed.2022.10.001

(3) Lanata, C. F.; Fischer-Walker, C. L.; Olascoaga, A. C.; Torres, C. X.; Aryee, M. J.; Black, R. E.; for the Child Health Epidemiology Reference Group of the World Health Organization and Unicef. Global Causes of Diarrheal Disease Mortality in Children <5 Years of Age: A Systematic Review. PLOS ONE 2013, 8 (9), e72788. 10.1371/journal.pone.0072788

(4) Baggen, J.; Thibaut, H. J.; Strating, J. R. P. M.; van Kuppeveld, F. J. M. The Life Cycle of Non-Polio Enteroviruses and How to Target It. Nature Reviews Microbiology 2018, 16 (6), 368–381. 10.1038/s41579-018-0005-4

(5) International Committee on Taxonomy of Viruses (ICTV). Genus: Enterovirus. ICTV Report. https://ictv.global/report/chapter/picornaviridae/picornaviridae/enterovirus (accessed 2025-10-12)

(6) International Committee on Taxonomy of Viruses (ICTV). Genus: Mastadenovirus. ICTV Report. https://ictv.global/report/chapter/adenoviridae/adenoviridae/mastadenovirus (accessed 2025-10-17)

(7) Liu, H.; Jin, L.; Koh, S. B. S.; Atanasov, I.; Schein, S.; Wu, L.; Hong Zhou, Z. Atomic Structure of Human Adenovirus by Cryo-EM Reveals Interactions Among Protein Networks. Science 2010, 329 (5995), 1038–1043. 10.1126/science.1187433

(8) van Raaij, M. J.; Louis, N.; Chroboczek, J.; Cusack, S. Structure of the Human Adenovirus Serotype 2 Fiber Head Domain at 1.5 Å Resolution. Virology 1999, 262 (2), 333–343. 10.1006/viro.1999.9849

(9) Lodder, W. J.; de Roda Husman, A. M. Presence of Noroviruses and Other Enteric Viruses in Sewage and Surface Waters in The Netherlands. Applied and Environmental Microbiology 2005, 71 (3), 1453–1461. 10.1128/AEM.71.3.1453-1461.2005

(10) Lodder, W. J.; van den Berg, H. H. J. L.; Rutjes, S. A.; de Roda Husman, A. M. Presence of Enteric Viruses in Source Waters for Drinking Water Production in the Netherlands. Applied and Environmental Microbiology 2010, 76 (17), 5965–5971. 10.1128/AEM.00245-10

(11) Wyn-Jones, A. P.; Carducci, A.; Cook, N.; D’Agostino, M.; Divizia, M.; Fleischer, J.; Gantzer, C.; Gawler, A.; Girones, R.; Höller, C.; de Roda Husman, A. M.; Kay, D.; Kozyra, I.; López-Pila, J.; Muscillo, M.; José Nascimento, M. S.; Papageorgiou, G.; Rutjes, S.; Sellwood, J.; Szewzyk, R.; Wyer, M. Surveillance of Adenoviruses and Noroviruses in European Recreational Waters. Water Research 2011, 45 (3), 1025–1038. 10.1016/j.watres.2010.10.015

(12) Vergara, G. G. R. V.; Rose, J. B.; Gin, K. Y. H. Risk Assessment of Noroviruses and Human Adenoviruses in Recreational Surface Waters. Water Research 2016, 103, 276–282. 10.1016/j.watres.2016.07.048

(13) Boehm, A. B.; Silverman, A. I.; Schriewer, A.; Goodwin, K. Systematic Review and Meta-Analysis of Decay Rates of Waterborne Mammalian Viruses and Coliphages in Surface Waters. Water Research 2019, 164, 114898. 10.1016/j.watres.2019.114898

(14) Ward, R. L.; Knowlton, D. R.; Winston, P. E. Mechanism of Inactivation of Enteric Viruses in Fresh Water. Applied and Environmental Microbiology 1986, 52 (3), 450–459. 10.1128/aem.52.3.450-459.1986

(15) Corre, M.-H.; Bachmann, V.; Kohn, T. Bacterial Matrix Metalloproteases and Serine Proteases Contribute to the Extra-Host Inactivation of Enteroviruses in Lake Water. The ISME Journal 2022, 16 (8), 1970–1979. 10.1038/s41396-022-01246-3

(16) Meibom, J.; Torii, S.; Morales, L. D.; Zumstein, M.; Kohn, T. A Capsid Tyrosine Residue Governs the Proteolytic Inactivation of Echovirus 11 in Lakewater. Environmental Science & Technology 2026. 10.1021/acs.est.6c08363

(17) Cliver, D. O.; Herrmann, J. E. Proteolytic and Microbial Inactivation of Enteroviruses. Water Research 1972, 6 (7), 797–805. 10.1016/0043-1354(72)90032-2

(18) Myouga, H.; Yoshimizu, M.; Tajima, K.; Ezura, Y. Purification of an Antiviral Substance Produced by Alteromonas Sp. and Its Virucidal Activity against Fish Viruses. Fish Pathology 1995, 30 (1), 15–22. 10.3147/jsfp.30.15

(19) Herrmann, J. E.; Cliver, D. O. Degradation of Coxsackievirus Type A9 by Proteolytic Enzymes. Infection and immunity 1973, 7 (4), 513–517. 10.1128/iai.7.4.513-517.1973

(20) Qin, Y.; Wang, J.; Wang, F.; Shen, L.; Zhou, H.; Sun, H.; Hao, K.; Song, L.; Zhou, Z.; Zhang, C.; Wu, Y.; Yang, J. Purification and Characterization of a Secretory Alkaline Metalloprotease with Highly Potent Antiviral Activity from *Serratia Marcescens* Strain S3. Journal of Agricultural and Food Chemistry 2019, 67 (11), 3168–3178. 10.1021/acs.jafc.8b06909

(21) Corre, M.-H.; Rey, B.; David, S. C.; Torii, S.; Chiappe, D.; Kohn, T. The Early Communication Stages between Serine Proteases and Enterovirus Capsids in the Race for Viral Disintegration. Communications Biology 2024, 7 (1), 969. 10.1038/s42003-024-06627-2

(22) Morvan, C.; Nekoua, M. P.; Mbani, C. J.; Debuysschere, C.; Alidjinou, E. K.; Hober, D. Enteroviruses in Water: Epidemiology, Detection and Inactivation. Environmental Microbiology 2025, 27 (5). 10.1111/1462-2920.70109

(23) Hot, D.; Legeay, O.; Jacques, J.; Gantzer, C.; Caudrelier, Y.; Guyard, K.; Lange, M.; Andréoletti, L. Detection of Somatic Phages, Infectious Enteroviruses and Enterovirus Genomes as Indicators of Human Enteric Viral Pollution in Surface Water. Water Research 2003, 37 (19), 4703–4710. 10.1016/S0043-1354(03)00439-1

(24) Tiemessen, C. T.; Kidd, A. H. Adenovirus Type 40 and 41 Growth in Vitro: Host Range Diversity Reflected by Differences in Patterns of DNA Replication. Journal of Virology 1994, 68 (2), 1239–1244. 10.1128/jvi.68.2.1239-1244.1994

(25) Carratalà, A.; Shim, H.; Zhong, Q.; Bachmann, V.; Jensen, J. D.; Kohn, T. Experimental Adaptation of Human Echovirus 11 to Ultraviolet Radiation Leads to Resistance to Disinfection and Ribavirin. Virus Evolution 2017, 3 (2), vex035. 10.1093/ve/vex035

(26) Ferguson, M.; Ihrie, J. MPN: Most Probable Number and Other Microbial Enumeration Techniques, 2024. https://cran.r-project.org/web/packages/MPN/index.html (accessed 2025-11-12)

(27) Hornung, R. W.; Reed, L. D. Estimation of Average Concentration in the Presence of Nondetectable Values. Applied Occupational and Environmental Hygiene 1990, 5 (1), 46–51. 10.1080/1047322X.1990.10389587

(28) Romanenko, A.; Peter, H.; Meibom, J.; Borchardt, M. A.; Kohn, T. Diversity of Lake Bacteria Promotes Human Echovirus Inactivation. Applied and Environmental Microbiology 2025, 91 (2), e02366–24. 10.1128/aem.02366-24

(29) Olive, M.; Gan, C.; Carratalà, A.; Kohn, T. Control of Waterborne Human Viruses by Indigenous Bacteria and Protists Is Influenced by Temperature, Virus Type, and Microbial Species. Applied and Environmental Microbiology 2020, 86 (3), e01992–19. 10.1128/AEM.01992-19

(30) Nuanualsuwan, S.; Cliver, D. O. Capsid Functions of Inactivated Human Picornaviruses and Feline Calicivirus. Applied and Environmental Microbiology 2003, 69 (1), 350–357. 10.1128/AEM.69.1.350-357.2003

(31) Schaub, A.; Luo, B.; David, S. C.; Glas, I.; Klein, L. K.; Costa, L.; Terrettaz, C.; Bluvshtein, N.; Motos, G.; Violaki, K.; Pohl, M. O.; Hugentobler, W.; Nenes, A.; Stertz, S.; Krieger, U. K.; Peter, T.; Kohn, T. Salt Supersaturation as an Accelerator of Influenza A Virus Inactivation in 1 _μ_L Droplets. Environmental Science & Technology 2024, 58 (42), 18856–18869. 10.1021/acs.est.4c04734

(32) Gall, A. M.; Shisler, J. L.; Mariñas, B. J. Analysis of the Viral Replication Cycle of Adenovirus Serotype 2 after Inactivation by Free Chlorine. Environ. Sci. Technol. 2015, 49 (7), 4584–4590. 10.1021/acs.est.5b00301

(33) Shin, S.; Lee, Y.; Kohn, T. Unraveling the Inactivation Mechanisms of Human Adenovirus 2 in Sunlight Disinfection: Synergism between Direct and Indirect Pathways. Environmental Science & Technology 2025, 59 (38), 20608–20615. 10.1021/acs.est.5c06149

(34) Rossmann, M. G.; He, Y.; Kuhn, R. J. Picornavirus–Receptor Interactions. Trends in Microbiology 2002, 10 (7), 324–331. 10.1016/S0966-842X(02)02383-1

(35) Wolf, C.; von Gunten, U.; Kohn, T. Kinetics of Inactivation of Waterborne Enteric Viruses by Ozone. Environmental Science & Technology 2018, 52 (4), 2170–2177. 10.1021/acs.est.7b05111

(36) Cromeans, T. L.; Kahler, A. M.; Hill, V. R. Inactivation of Adenoviruses, Enteroviruses, and Murine Norovirus in Water by Free Chlorine and Monochloramine. Applied and Environmental Microbiology 2010, 76 (4), 1028–1033. 10.1128/AEM.01342-09

(37) Torii, S.; Corre, M.-H.; Miura, F.; Itamochi, M.; Haga, K.; Katayama, K.; Katayama, H.; Kohn, T. Genotype-Dependent Kinetics of Enterovirus Inactivation by Free Chlorine and Ultraviolet (UV) Irradiation. Water Research 2022, 220, 118712. 10.1016/j.watres.2022.118712

(38) Wigginton, K. R.; Pecson, B. M.; Sigstam, T.; Bosshard, F.; Kohn, T. Virus Inactivation Mechanisms: Impact of Disinfectants on Virus Function and Structural Integrity. Environmental Science & Technology 2012, 46 (21), 12069–12078. 10.1021/es3029473

(39) Meibom, J.; Wichmann, N.; Astorch-Cardona, A.; Zumstein, M.; Kohn, T. Proteolytic Activity and Substrate Specificity of Lake Geneva. Environmental Science & Technology 2025, 59 (51), 27811–27823. 10.1021/acs.est.5c06582

(40) Yamamoto, S.; Ogasawara, N.; Sudo-Yokoyama, Y.; Sato, S.; Takata, N.; Yokota, N.; Nakano, T.; Hayashi, K.; Takasawa, A.; Endo, M.; Hinatsu, M.; Yoshida, K.; Sato, T.; Takahashi, S.; Takano, K.; Kojima, T.; Hiraki, J.; Yokota, S. I. *Bacillaceae* Serine Proteases and *Streptomyces* Epsilon-Poly-l-Lysine Synergistically Inactivate *Caliciviridae* by Inhibiting RNA Genome Release. Scientific Reports 2024, 14 (1), 15181. 10.1038/s41598-024-65963-9

(41) Kimura, T.; Yoshimizu, M.; Ezura, Y.; Kamei, Y. An Antiviral Agent (46NW-04A) Produced by *Pseudomonas* Sp. and Its Activity against Fish Viruses. Journal of Aquatic Animal Health 1990, 2 (1), 12–20. 10.1577/1548-8667(1990)002%253C0012:AAAPBP%253E2.3.CO;2

(42) Padhi, C.; Field, C. M.; Forneris, C. C.; Olszewski, D.; Fraley, A. E.; Sandu, I.; Scott, T. A.; Farnung, J.; Ruscheweyh, H.-J.; Narayan Panda, A.; Oxenius, A.; Greber, U. F.; Bode, J. W.; Sunagawa, S.; Raina, V.; Suar, M.; Piel, J. Metagenomic Study of Lake Microbial Mats Reveals Protease-Inhibiting Antiviral Peptides from a Core Microbiome Member. Proceedings of the National Academy of Sciences 2024, 121 (49), e2409026121. 10.1073/pnas.2409026121

(43) Bello-Morales, R.; Andreu, S.; Ruiz-Carpio, V.; Ripa, I.; López-Guerrero, J. A. Extracellular Polymeric Substances: Still Promising Antivirals. Viruses 2022, 14 (6), 1337. 10.3390/v14061337

(44) Kuznetsova, T. A.; Besednova, N. N.; Zaporozhets, T. S.; Kokoulin, M. S.; Khotimchenko, Y. S.; Shchelkanov, M. Y. Antiviral Potential of Marine Bacteria Polysaccharides. Russian Journal of Marine Biology 2024, 50 (3), 107–115. 10.1134/S1063074024700056

(45) Wang, X.; Hu, W.; Zhu, L.; Yang, Q. *Bacillus Subtilis* and Surfactin Inhibit the Transmissible Gastroenteritis Virus from Entering the Intestinal Epithelial Cells. Bioscience Reports 2017, 37 (2), BSR20170082. 10.1042/BSR20170082

(46) Gelbart, W. M.; Knobler, C. M. Pressurized Viruses. Science 2009, 323 (5922), 1682–1683. 10.1126/science.1170645

(47) Pérez-Berná, A. J.; Ortega-Esteban, A.; Menéndez-Conejero, R.; Winkler, D. C.; Menéndez, M.; Steven, A. C.; Flint, S. J.; de Pablo, P. J.; San Martín, C. The Role of Capsid Maturation on Adenovirus Priming for Sequential Uncoating. Journal of Biological Chemistry 2012, 287 (37), 31582–31595. 10.1074/jbc.M112.389957

(48) Kuijpers, L.; Giannopoulou, E.-A.; Feng, Y.; van den Braak, W.; Freydoonian, A.; Ramlal, R.; Meiring, H.; Solano, B.; Roos, W. H.; Jakobi, A. J.; van der Pol, L. A.; Dekker, N. H. Enterovirus-like Particles Encapsidate RNA and Exhibit Decreased Stability Due to Lack of Maturation. PLoS Pathogens 2025, 21 (2), e1012873. 10.1371/journal.ppat.1012873

(49) Zubieta, C.; Schoehn, G.; Chroboczek, J.; Cusack, S. The Structure of the Human Adenovirus 2 Penton. Molecular Cell 2005, 17 (1), 121–135. 10.1016/j.molcel.2004.11.041

(50) Thurston-Enriquez, J. A.; Haas, C. N.; Jacangelo, J.; Gerba, C. P. Chlorine Inactivation of Adenovirus Type 40 and Feline Calicivirus. Applied and Environmental Microbiology 2003, 69 (7), 3979–3985. 10.1128/AEM.69.7.3979-3985.2003

(51) Kahler, A. M.; Cromeans, T. L.; Metcalfe, M. G.; Humphrey, C. D.; Hill, V. R. Aggregation of Adenovirus 2 in Source Water and Impacts on Disinfection by Chlorine. Food and Environmental Virology 2016, 8 (2), 148–155. 10.1007/s12560-016-9232-x

(52) Thurston-Enriquez, J. A.; Haas, C. N.; Jacangelo, J.; Gerba, C. P. Inactivation of Enteric Adenovirus and Feline Calicivirus by Chlorine Dioxide. Applied and Environmental Microbiology 2005, 71 (6), 3100–3105. 10.1128/AEM.71.6.3100-3105.2005

(53) Sigstam, T.; Rohatschek, A.; Zhong, Q.; Brennecke, M.; Kohn, T. On the Cause of the Tailing Phenomenon during Virus Disinfection by Chlorine Dioxide. Water Research 2014, 48, 82–89. 10.1016/j.watres.2013.09.023

(54) Lytle, C. D.; Sagripanti, J.-L. Predicted Inactivation of Viruses of Relevance to Biodefense by Solar Radiation. Journal of Virology 2005, 79 (22), 14244–14252. 10.1128/jvi.79.22.14244-14252.2005

(55) Meister, S.; Verbyla, M. E.; Klinger, M.; Kohn, T. Variability in Disinfection Resistance between Currently Circulating *Enterovirus B* Serotypes and Strains. Environmental Science & Technology 2018, 52 (6), 3696–3705. 10.1021/acs.est.8b00851

(56) Verbel-Olarte, M. I.; Kohn, T.; Ismail, N. S. Differential Effects of Zooplankton on Sunlight Inactivation of Viruses. bioRxiv March 5, 2026, p 2026.03.05.709857. 10.64898/2026.03.05.709857

