## Supplementary material for "Distinct kinetics and mechanisms of microbial inactivation of enteric virus revealed by capsid and genome Integrity": Meibom-et-al 2026 Supplemental

Number of pages: 10

Number of tables: 3

Number of figures: 6

Supplementary Methods

*Virus size measurement by dynamic light scattering (DLS)*

Hydrodynamic size measurements of E11, CVB5, and HAdV2 virus stocks (**Supplementary Figure S1**) were performed using the Zetasizer Nano ZS (Malvern Instruments). Briefly, 400 μL virus stocks (diluted 1:40 in PBS) were added to 1.5 mL disposable cuvettes (759150, Brand Gmbh). The cuvettes were placed in the Zetasizer Nano ZS instrument taking care to always maintain the same orientation. Data acquisition was performed with the Zetasizer Software (v6.12, Malvern Instruments) using backscattering mode (173°). The absorption and refractive index (RI) were set to 0.01 and 1.590 (RI of polysterene latex), respectively. The temperature and viscosity of the dispersant were set to 22°C and 0.9540 mPaˑs, respectively. Three runs of 10 seconds were averaged per measurement.

*Enumeration of virus genome copies: (RT-)dPCR details*

RT-dPCR for E11 and CVB5 was performed using the QIAcuity OneStep Advanced Probe Kit (250132, Qiagen) with RNA extracts (undiluted or diluted 1:1000) as templates and the primer pair and probe listed in **Supplementary** **Table S1**. The reaction volumes were 12 μL and consisted each of 3 μL 4× OneStep Advanced Probe Master Mix, 0.12 μL OneStep RT Mix, 1.5 μL Enhancer GC, 0.6 μL each of forward and reverse primers (20 μM), 0.3 μL probe (20 μM), 4 μL RNA template, and 1.88 μL nuclease-free water. RT-dPCR was performed on the QIAcuity One 2-plex Device (Qiagen) using 8.5k 96-well Nanoplates (250021, Qiagen) with the cycling parameters listed in **Supplementary** **Table S2**.

dPCR for HAdV2, was performed using the QIAcuity Probe PCR Kit (250101, Qiagen) with DNA extracts (diluted 1:10) as templates and the primer pair and probe listed in **Supplementary** **Table S1**. The reaction volumes were 12 μL and consisted each of 3 μL 4x QIAcuity Probe PCR Master Mix, 0.2 μL MluI-HF restriction enzyme (R3198S, New England Biolabs), 1.2 μL each of forward and reverse primers (10 μM), 0.6 μL probe (10 μM), 4 μL DNA template, and 1.8 μL nuclease-free water. The reaction mixtures were incubated for 10 minutes at room temperature prior to dPCR analysis to allow genome cleavage by the restriction enzyme. dPCR was performed on the QIAcuity One 2-plex Device (Qiagen) using 8.5k 96-well Nanoplates (250021, Qiagen) with the cycling parameters listed in **Supplementary** **Table S2**.

In each dPCR run, a negative control (nuclease-free water) and a positive control (virus stock genome extract diluted 1:100) were included. The primers and probes were purchased from Microsynth, Switzerland. Results were analyzed using QIAcuity Software Suite v2.1.7.182 (911001, Qiagen) with automatic settings for positive and negative partition separation.

*Direct assessment of HAdV2 fiber protein function: long-range PCR followed by qPCR details*

Long-range PCR targeting the adenovirus hexon gene was performed with undiluted DNA extracts as templates and the primer pair and cycling parameters listed in **Supplementary** **Tables S1 and S2**, respectively. The reaction volumes were 50 μL and consisted each of 10 μL 5x colorless GoTaq buffer (M7401, Promega), 0.25 μL GoTaq G2 Hot Start Polymerase (M7401, Promega), 4.75 μL MgCl_2_ (25 mM, M7401, Promega), 1 μL dNTP Mix (10 mM, U1511, Promega), 12 μL each of forward and reverse primers (4.2 μM), and 10 μL DNA template. Following PCR amplification, samples were treated with S1 Nuclease (M5761, Promega) for 30 minutes at 37°C and subsequently purified using the QIAquick PCR & Gel Cleanup Kit (28506, Qiagen) according to the manufacturer’s instructions.

qPCR was performed using purified PCR amplicons (diluted 1:10’000) as DNA templates, the primer pair and probe listed in **Supplementary** **Table S1**, and the cycling parameters listed in **Supplementary** **Table S2**. The reaction volumes were 25 μL and consisted each of 12.5 μL 2x TaqMan Environmental Master Mix (4396838, Applied Biosystems), 1 μL each of forward and reverse primers (22.5 μM), 0.5 μL probe (11.25 μM), and 10 μL DNA template.

Supplementary Tables

**Supplementary Table S1**. List of primers and probes used for quantification of viral genome copies by digital PCR.

| Virus | Primer | 5'-Sequence-3' |
| --- | --- | --- |
| E11 & CVB5^1^ | Forward | CCTCCGGCCCCTGAAT |
|  | Reverse | ACCGGATGGCCAATCCAA |
|  | Probe | HEX-CGGAACCGACTACTTTGGGTGTCCGT-BHQ1 |
| HAdV2^2,3^ | Forward | CWTACATGCACATCKCSGG |
|  | Reverse | CRCGGGCRAAYTGCACCAG |
|  | Probe | 6-FAM-CCGGGCTCAGGTACTCCGAGGCGTCCT-BHQ1 |
| HAdV2 fiber protein– long-range PCR^4^ | Forward | CACGGAGAGATGGCTATGCGCGGCGGTATCCTGCCCCTCC |
|  | Reverse | CGTAGGTGC ACCGTGGGGTTTCTAAAC |
| HAdV2 fiber protein–qPCR^4^ | Forward | CACGGAGAGATGGCTATGCG |
|  | Reverse | CAAGCGAGCGTGAGACTCC |
|  | Probe | FAM-TTGCATCCGTGGCCTTGCAGGCGCA-BHQ1 |

**Supplementary Table S2**. Cycling parameters for quantification of viral genome copies by digital PCR.

| Virus | RT step | Enzyme activation | Amplification | Final extension |
| --- | --- | --- | --- | --- |
| E11 & CVB5 | 50°C 40 min | 95°C 2min | 45 cycles: 95°C 15 sec, 60°C 1 min |  |
| HAdV2 |  | 95°C 2min | 50 cycles: 95°C 15 sec, 57°C 1 min |  |
| HAdV2 fiber protein– long-range PCR |  | 95°C 2 min | 35 cycles: 95°C 15 sec, 65°C 20 sec, 72°C 90 sec | 72°C 5 min |
| HAdV2 fiber protein–qPCR |  | 95°C 10 min | 40 cycles: 95°C 15 sec, 60°C 1 min |  |

**Supplementary Table S3**. Inactivation rate constants (*k*) of E11, CVB5, and HAdV2 in microbially active Lake Geneva surface water collected in April 2024, July 2024, and March 2026. Rate constants were determined in the linear range of inactivation, corresponding to the entire timeseries (0 – 54 hours) for all samples except those indicated by an asterisk (*) where the linear range was 0 – 30 hours. Data points below the LOQ were not considered for the calculation of rate constants.

| Sample | Virus | *k* ± 95% CI [h^-1^] |
| --- | --- | --- |
| April 2024 | E11 | 0.11 ± 0.01 |
|  | CVB5 | 0.02 ± 0.01 |
|  | HAdV2 * | 0.29 ± 0.03 |
| July 2024 | E11 | 0.09 ± 0.01 |
|  | CVB5 | 0.12 ± 0.01 |
|  | HAdV2 | 0.10 ± 0.01 |
| March 2026 | E11 | 0.10 ± 0.01 |
|  | CVB5 | 0.00 ± 0.00 |
|  | HAdV2 * | 0.25 ± 0.02 |

Supplementary Figures

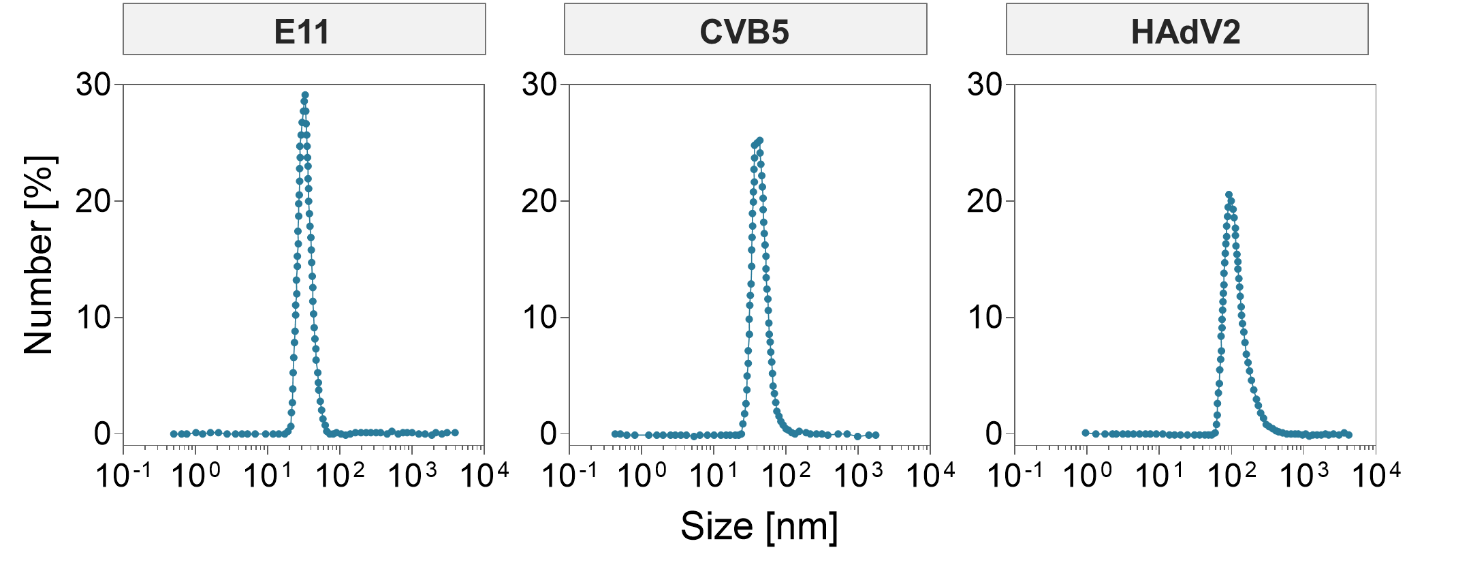

**Supplementary Figure S1**. Size distribution of E11, CVB5, and HAdV2 stocks in PBS measured by dynamic light scattering (DLS). Each measurement represents the average of three runs of 30 seconds.

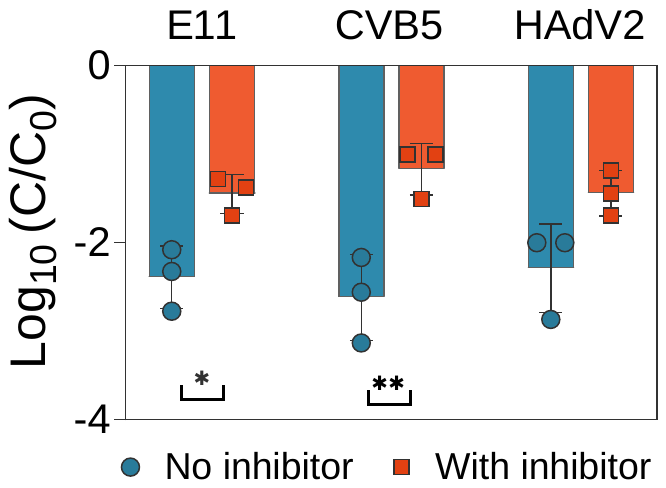

**Supplementary Figure S2**. Inactivation of E11, CVB5, and HAdV2 (t = 54 hours) in Lake Geneva surface water in the absence (blue circles) or presence (red squares) of the metalloprotease inhibitor GM6001. Lakewater water was collected in July 2024. Quantitative results of triplicate experiments are presented as individual data points with the mean and standard deviation indicated by the bar and error bars, respectively. Significant differences between Log_10_ inactivation were determined by paired t-test (*: p < 0.1; **: p < 0.01). Data for E11 has been previously reported^5^.

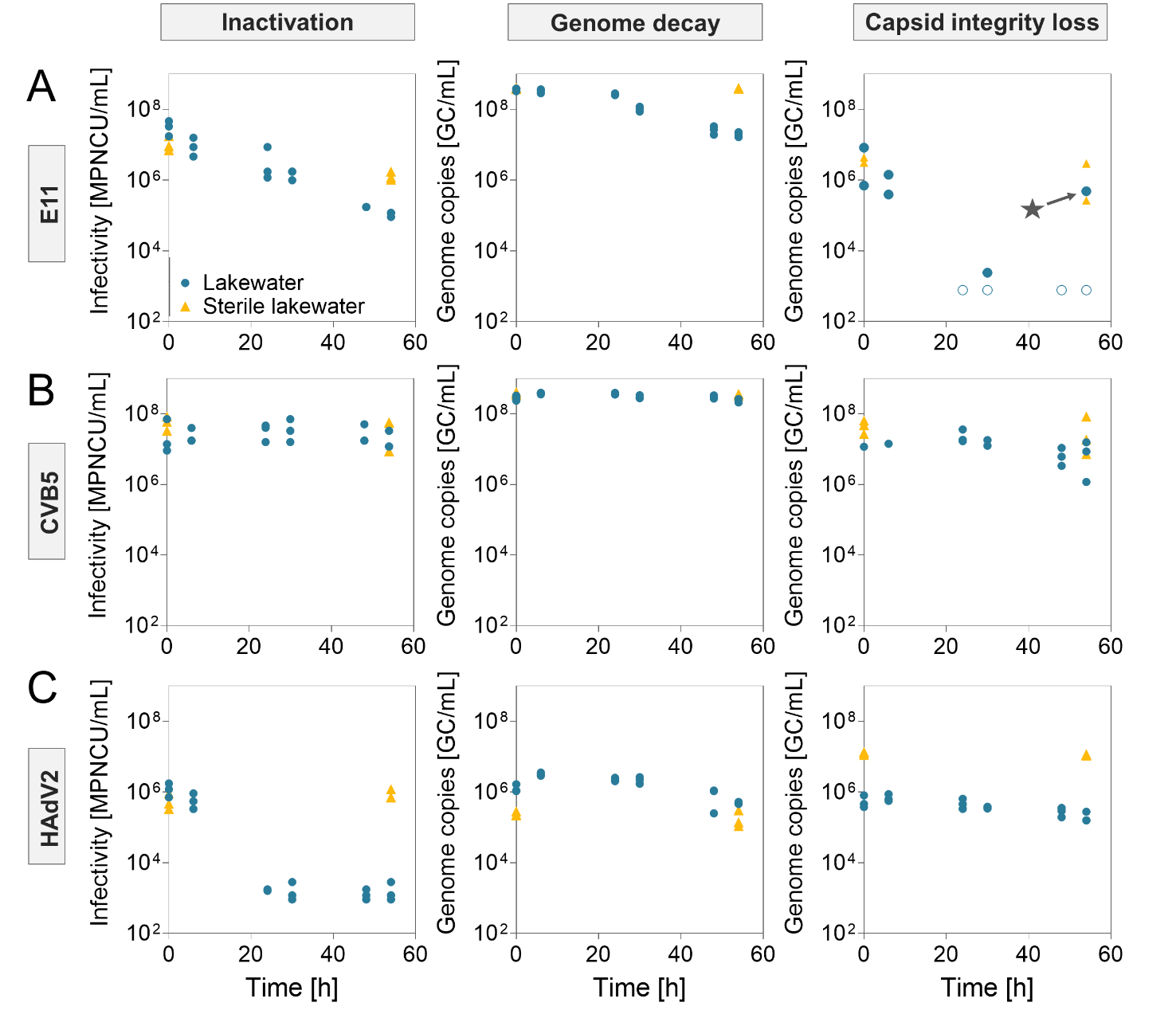

**Supplementary Figure S3.** Measured infectious virus particles (MPN) and viral genome copies (GC) of **(A)** E11, **(B)** CVB5, and **(C)** HAdV2 in microbially active (blue circles) and sterile (orange triangles) Lake Geneva surface water collected in March 2026. Virus genome copies were quantified before (middle column) or after (right column) nuclease treatment. Individual data points of triplicate experiments are presented, except for nuclease-treated E11 and CVB5 samples where two and one replicates are presented, respectively, for some timepoints. Data points below the dPCR detection limit were set to a concentration of LOQ/√2, with the corresponding Log_10_ genome decay indicated by an open symbol. The data point indicated by a star is an outlier and indicates likely contamination.

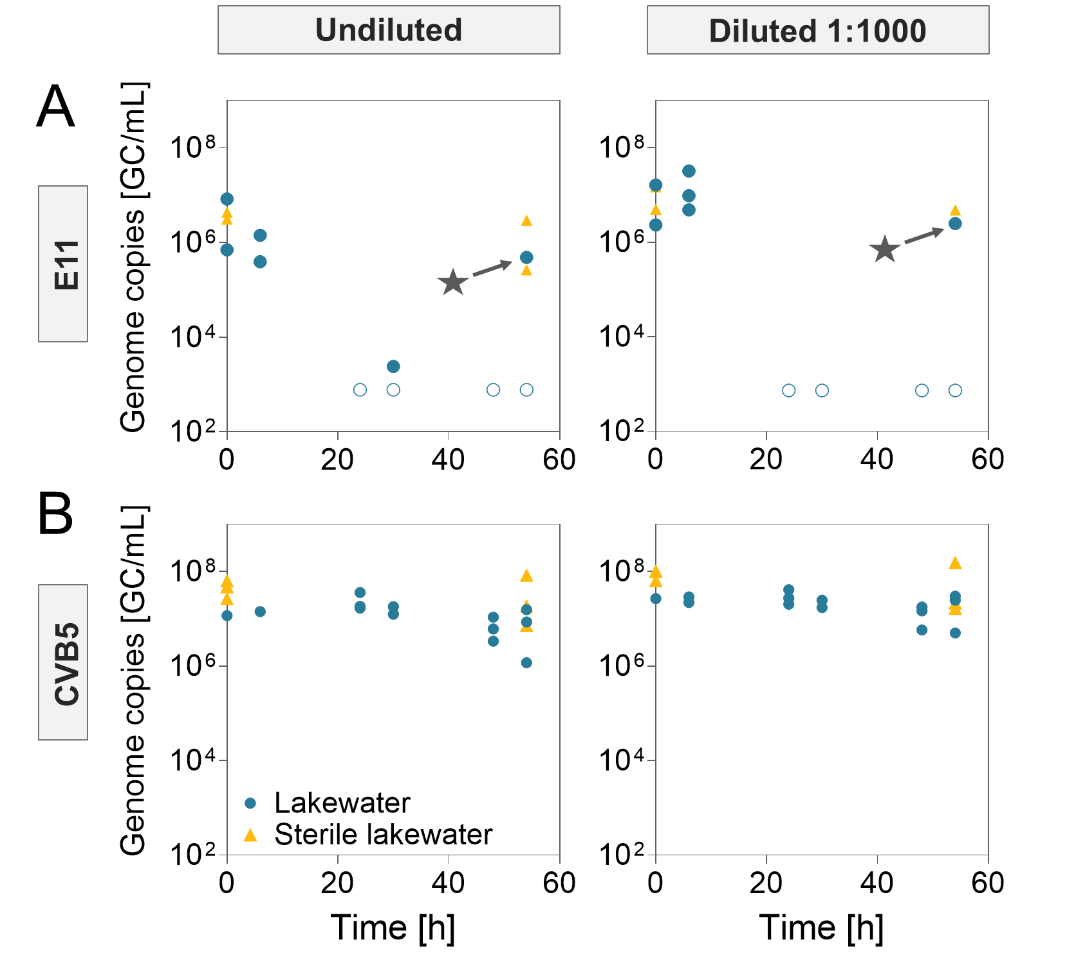

**Supplementary Figure S4.** Measured genome copies (GC) of **(A)** E11 and **(B)** CVB5 in microbially active (blue circles) and sterile (orange triangles) Lake Geneva surface water collected in March 2026. Virus genome copies were quantified after nuclease treatment, using either undiluted (left column) or 1000-fold diluted (right column) RNA extracts as dPCR templates. Data points below the dPCR detection limit were set to a concentration of LOQ/√2, with the corresponding Log_10_ genome decay indicated by an open symbol. The data points indicated by a star are outliers and indicates likely contamination.

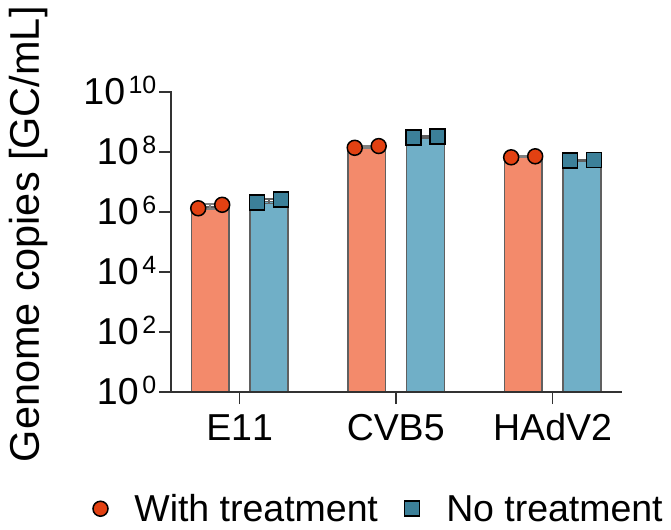

**Supplementary Figure S5.** Measured genome copies (GC) of E11, CVB5, and HAdV2 in microbially active Lake Geneva surface water (t = 0 hours) collected in November 2025. Virus genome copies were quantified before (blue squares) or after (red circles) nuclease treatment. Individual data points of duplicate experiments are presented.

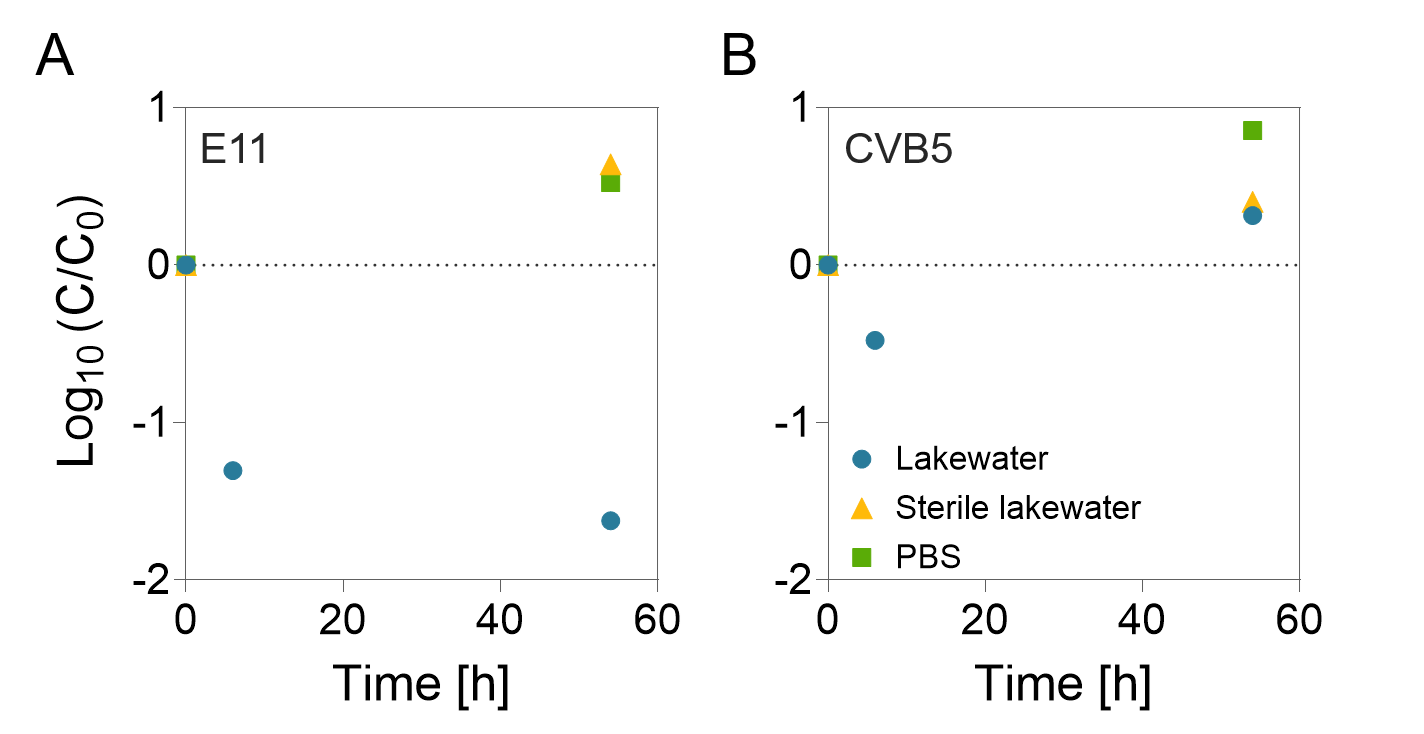

**Supplementary Figure S6.** Inactivation of **(A)** E11 and **(B)** CVB5 in microbially active lakewater (blue circles), sterile lakewater (orange triangles), and phosphate-buffered saline (PBS, green squares) performed in parallel to the Western blot analysis of viral protein 1 (VP1, **Figure 4**). Lake Geneva surface water was collected in March 2026. n = 1.

References

(1)	Romanenko, A.; Peter, H.; Meibom, J.; Borchardt, M. A.; Kohn, T. Diversity of Lake Bacteria Promotes Human Echovirus Inactivation. *Applied and Environmental Microbiology* **2025**, *91* (2), e02366-24. https://doi.org/10.1128/aem.02366-24

(2)	Hernroth, B. E.; Conden-Hansson, A.-C.; Rehnstam-Holm, A.-S.; Girones, R.; Allard, A. K. Environmental Factors Influencing Human Viral Pathogens and Their Potential Indicator Organisms in the Blue Mussel, *Mytilus Edulis*: The First Scandinavian Report. *Applied and Environmental Microbiology* **2002**, *68* (9), 4523–4533. https://doi.org/10.1128/AEM.68.9.4523-4533.2002

(3)	Bofill-Mas, S.; Albinana-Gimenez, N.; Clemente-Casares, P.; Hundesa, A.; Rodriguez-Manzano, J.; Allard, A.; Calvo, M.; Girones, R. Quantification and Stability of Human Adenoviruses and Polyomavirus JCPyV in Wastewater Matrices. *Applied and Environmental Microbiology* **2006**, *72* (12), 7894–7896. https://doi.org/10.1128/AEM.00965-06

(4)	Shin, S.; Lee, Y.; Kohn, T. Unraveling the Inactivation Mechanisms of Human Adenovirus 2 in Sunlight Disinfection: Synergism between Direct and Indirect Pathways. *Environmental Science & Technology* **2025**, *59* (38), 20608–20615. https://doi.org/10.1021/acs.est.5c06149

(5)	Meibom, J.; Torii, S.; Morales, L. D.; Zumstein, M.; Kohn, T. A Capsid Tyrosine Residue Governs the Proteolytic Inactivation of Echovirus 11 in Lakewater. *Environmental Science & Technology* **2026**. https://doi.org/10.1021/acs.est.6c08363
